# The Dynamics of Plant Nutation

**DOI:** 10.1101/2020.05.11.089466

**Authors:** Vicente Raja, Paula L. Silva, Roghaieh Holghoomi, Paco Calvo

## Abstract

In this article we advance a cutting-edge methodology for the study of the dynamics of plant movements of nutation. Our approach, unlike customary kinematic analyses of shape, period, or amplitude, is based on three typical signatures of adaptively controlled processes and motions, as reported in the biological and behavioral dynamics literature: harmonicity, predictability, and complexity. We illustrate the application of a dynamical methodology to the bending movements of shoots of common beans (*Phaseolus vulgaris* L.) in two conditions: with and without a support to climb onto. The results herewith reported support the hypothesis that patterns of nutation are influenced by the presence of a support to climb in their vicinity. The methodology is in principle applicable to a whole range of plant movements.

---

Since first described extensively by Charles Darwin^[1, 2]^, bending movements of nutation have been studied in monocotyledons and dicotyledons^[3]^, in fungi, algae, or bryophytes^[4, 5]^, and in shoots and roots of climbing plants^[6 – 10]^, among others^[40 – 43]^. Most kinematic aspects of nutation in different plants species have been thoroughly researched—e.g., oscillatory shapes and directions^[11 - 13]^, period^[14, 15]^, or amplitude^[16, 17]^. Nutation kinematics of different organs has served to lay a foundation of several mechanisms postulated as responsible for the movement in question: internal oscillators^[2, 16, 18]^, gravitation-driven mechanisms^[19, 20]^, or a combination of these two mechanisms^[21 – 23]^. Moreover, nutation has been taken to be an instance of plants’ adaptively controlled motion^[2, 23 – 28]^. However, which one of those proposed mechanisms better accounts for the complex kinematics of plant nutation, both generally and in terms of adaptive control, requires experimental adjudication.

Different models inspired by very different assumptions aim to do so—assumptions that range from a purely nastic, non-directional nature of the movement^[13]^ to a full-fledged tropic rendering, including models that resort to kinematical landmarks deployed in the animal literature to characterize directional grasping behavior^[64]^. All such models nonetheless run the risk of generating the same average patterns—quasi-circular and quasi-elliptical patterns depending on their parametrization—despite those different assumptions^[13, 18, 19, 29 – 31]^. For this reason, current methods based solely on the kinematics of nutation are not able to decide between the different hypotheses for the control strategy and underlying mechanism of the movement, and its tropic and/or nastic components. A fundamental step to experimentally discern about these issues is to achieve a more profound understanding of the nature of plant nutation.

The aim of this article is to go beyond basic plant kinematics to improve our understanding of the nature and control strategies of plant nutation through a careful characterization of its underlying dynamical organization. To overcome the problems of the approaches based on kinematical patterns, we looked for methods described outside the field of plant science to capture the dynamical processes that give rise to those patterns. These methods, thoroughly used in the behavioral and biological sciences^[32 – 35]^, are not based on summary statistics or average patterns, blind to the temporal dependencies in the observed movement. Instead, the methods herewith reported are sensitive to such temporal dependencies, offering better access to the underlying dynamical organization that gives rise to them.

Our methodology is based on the analysis of time series gathered by a common procedure in the field—time-lapse, zenithal point-view recording of nutation^[7, 23, 27, 37]^—and provides measurements for some typical signatures of adaptively controlled motions reported in the literature on behavior dynamics: harmonicity, predictability, and complexity^[36, 38, 39, 46]^. The relationship between these signatures and adaptive controlled motions has to do with the temporal dependencies of these motions. As opposed to completely non-adaptive random motions, the states of adaptive controlled motions at any moment of time, although variable, are partially determined by their own previous states and by the environmental factors that influence them. The influences of those environmental factors make adaptive controlled motions more predictable with respect to them: they tend to re-visit similar locations due to those influences. And the combination of variability (randomness) and determinism is the signature of complex processes^[44, 53, 60, 68 – 71]^.This is the sense in which the proposed measurements are adequate to study adaptive controlled motions.

To illustrate the application of the methodology in the plant sciences, we compared the nutation dynamics of bean plants growing with and without a support (a pole) to twine around in otherwise identical environments. We hypothesize that if nutation is an adaptively controlled motion, the plants’ bending movements will exhibit a nonlinear organization that is more predictable and complex when the pole is present. We show how the general methodology is applied in this study and the way in which the measurement of the dynamical properties of nutation patterns speak to the control mechanism of the movement and the influence of the climbing support. Overall, our aim is twofold: (i) to illustrate the use of nonlinear methods to test hypothesis regarding the nature of plant nutation and the environmental factors that might influence it; and to (ii) furnish plant scientists with a set of guidelines for the processing of movements of nutation in a reliable and informative manner.

## Results

### Characterization of Nutation Data

To study the dynamical features of nutation patterns we designed an experimental analysis on the common bean (*Phaseolus vulgaris* L. var. Bueno Aires) based on established protocols^[23]^. Concretely, we recorded the behavior of twenty potted common bean plants in two different conditions: with and without a climbing support (a pole) in their vicinity. The twenty plants were placed pairwise in two identical recording cylindrical booths, one pair at a time (Figure 1). In one of the cabins, a pole was placed 30 centimeters away from the plant. There was no climbing support in the control recording cabin (see Methods).

**Figure 1.**
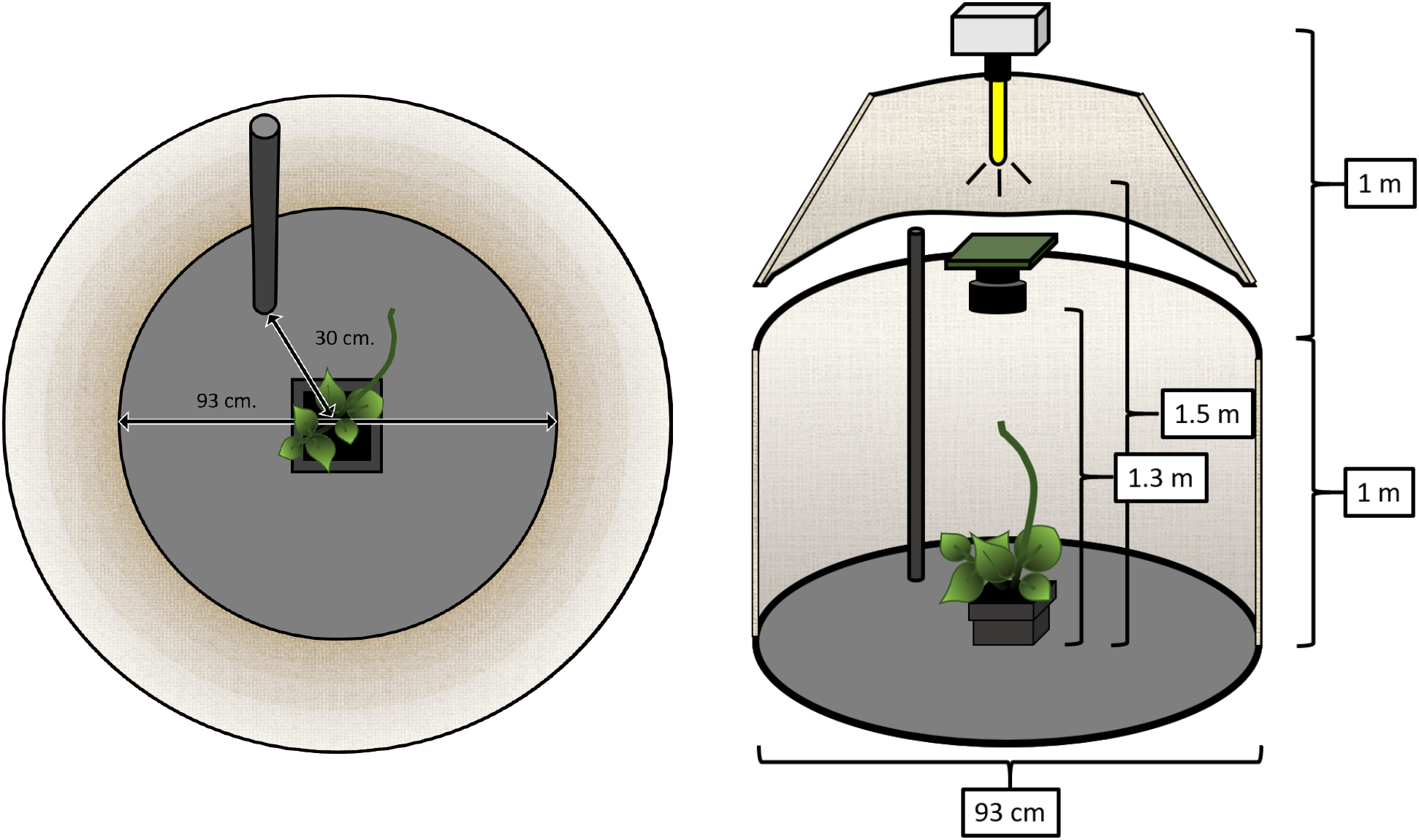
Diagram of the experimental setting (see Method for full details). Cylindrical booth for the pole condition. The cylindrical booth for the no-pole condition was identical except for the presence of the pole. ***Left:*** Zenithal camera viewpoint of a bean plant and the pole (30 centimeters away from the bean plant; height: 0.90 cm, diameter: 1.8 cm) in the cylindrical boot (width: 93 cm). ***Right:*** Lateral viewpoint of a bean plant and the pole in the cylindrical booth. *Time Lapse Camera:* Brinno TLC200 PRO (height: 130 cm); 4.2 μm High Dynamic Range (115dB) image sensor for recording in darkness. *Illumination:* high-pressure sodium lamp Lumatek pulse-start HPS Lamp 250W (height: 150 cm; photon fluence rate: 430 ± 50 μmol m–2 s–1 at leaf level). A white parabolic reflector provided symmetrical lighting.

Zenithal pictures were taken at one-minute intervals from the onset of plant nutation until the shoot tip of the bean made contact with the pole. Time-lapse footage was assembled out of the zenithal frames being taken (number of frames ranged from 2300 to 5300; M ± SD = 3576.5 ± 842.83). Using the *Circumnutation Tracker (CT)* software^[37]^, the position of each plant’s shoot tip in the horizontal plane was digitalized at a 6Hz rate and time series of the movement that ranged between 451 and 1049 data points (M ± SD = 715.3 ± 168.56) were generated (Figure 2B). Fast-Fourier transformations showed that most of the frequency power of the movement was concentrated under 0.5Hz (M ± SD = 0.29 ± 0.05; Figure 2A). To minimize high frequency noise, a low-pass filter with a cut-off frequency of 1.5Hz was used.

**Figure 2.**
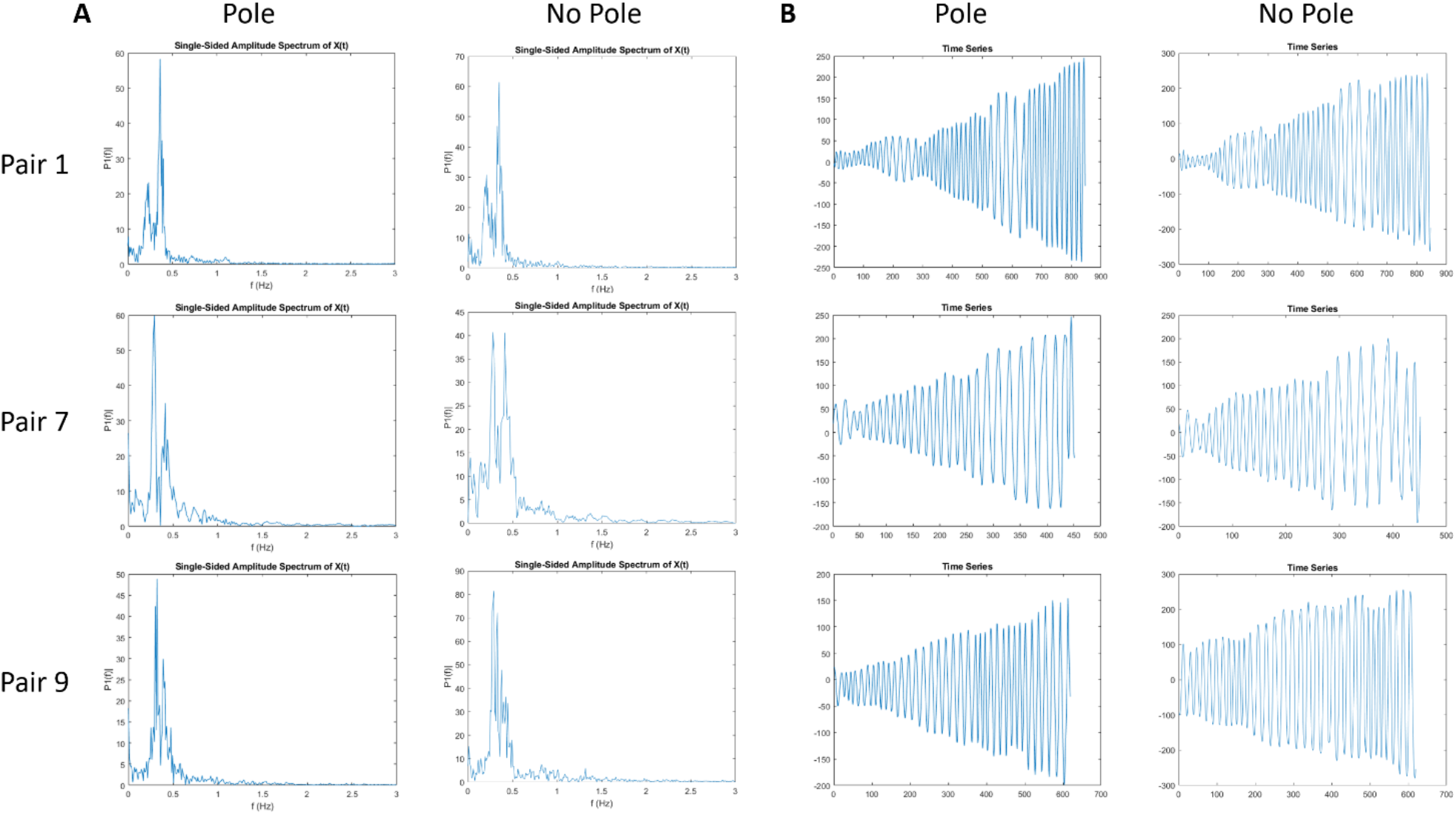
Three examples of the detail of the time series (Pair 1, Pair 7, and Pair 9). Every row corresponds to one pair of plants. **(A)** Fast-Fourier Transformation of the pairs of time series organized by condition. Main frequency components were around 0.15-0.35 Hz for all plants. **(B)** Time-series organized by condition. Displacement of the shoot-tip across the x-axis of the frames. Amplitude of the displacement of the shoot-tip (in pixels) is plotted against time (in samples).

To examine the effect of time on nutation dynamics, we divided the data into as many hoping windows of 100 data points as possible, starting from the last point defined by the moment at which the plant in the pole condition touched the pole. For most plants, this procedure guaranteed at least 6 windows of 100 points that were used for analysis. Amplitude and frequency of nutation changed over time, but detailed analysis showed that data was stationary within each window. Measures of harmonicity, predictability, and complexity of nutation patterns were computed for each plant in each of the six analyzed windows (Windows coded 1 to 6; see Table 1).

**Table 1.** Window selection for each pair of plants. Hoping windows (100 samples each) are labelled starting from the last window (6) and going backwards from it. The process of selection ensures homogeneity and stationarity of the data in the windows relevant for the analysis (Windows 1 to 6).

| Pair | Total | Samples per Window (Range) |  |  |  |  |  |  |  |  |  |  |
| --- | --- | --- | --- | --- | --- | --- | --- | --- | --- | --- | --- | --- |
|  |  | -4 | -3 | -2 | -1 | 0 | 1 | 2 | 3 | 4 | 5 | 6 |
| 1 | 846 | - | - | 0-45 | 46-145 | 146-245 | 246-345 | 346-445 | 446-545 | 546-645 | 646-745 | 746-846 |
| 2 | 1049 | 0-48 | 49-148 | 149-248 | 249-348 | 349-448 | 449-548 | 549-648 | 649-748 | 749-848 | 849-948 | 949-1049 |
| 3 | 795 | - | - | - | 0-94 | 95-194 | 195-294 | 295-394 | 395-494 | 495-594 | 595-694 | 695-795 |
| 4 | 760 | - | - | - | 0-59 | 60-159 | 160-259 | 260-359 | 360-459 | 460-559 | 560-659 | 660-760 |
| 5 | 655 | - | - | - | - | 0-54 | 55-154 | 155-254 | 255-354 | 355-454 | 455-554 | 555-655 |
| 6 | 750 | - | - | - | 0-49 | 50-149 | 150-249 | 250-349 | 360-449 | 450-549 | 550-649 | 650-750 |
| 7 | 451 | - | - | - | - | - | - | 0-50 | 51-150 | 151-250 | 251-350 | 351-451 |
| 8 | 528 | - | - | - | - | - | 0-27 | 28-127 | 128-227 | 228-327 | 328-427 | 428-528 |
| 9 | 619 | - | - | - | - | 0-18 | 19-118 | 119-218 | 219-318 | 319-418 | 419-518 | 519-619 |
| 10 | 699 | - | - | - | - | 0-98 | 99-198 | 199-298 | 299-398 | 399-498 | 499-598 | 599-699 |

### Harmonicity of Nutation Patterns: Normalized Peak Acceleration

Normalized Peak Acceleration (NPA) was computed to assess the degree of deviation of the recorded plant movements from a purely sinusoidal (harmonic) pattern. *Aω*^2^ corresponds to the peak acceleration of a sinusoidal movement of amplitude *A* and frequency *ω*^2^. Thus, a purely sinusoidal movement governed by linear dynamics gives rise to an NPA value of 1. Figure 3A displays the mean (and standard error) of NPA over time for the two conditions. Observed NPA values are larger than 1 and therefore indicate that nutation patterns are not purely sinusoidal or harmonic. These deviations from harmonicity in turn signal the contribution of non-linear processes to nutation.

**Figure 3.**
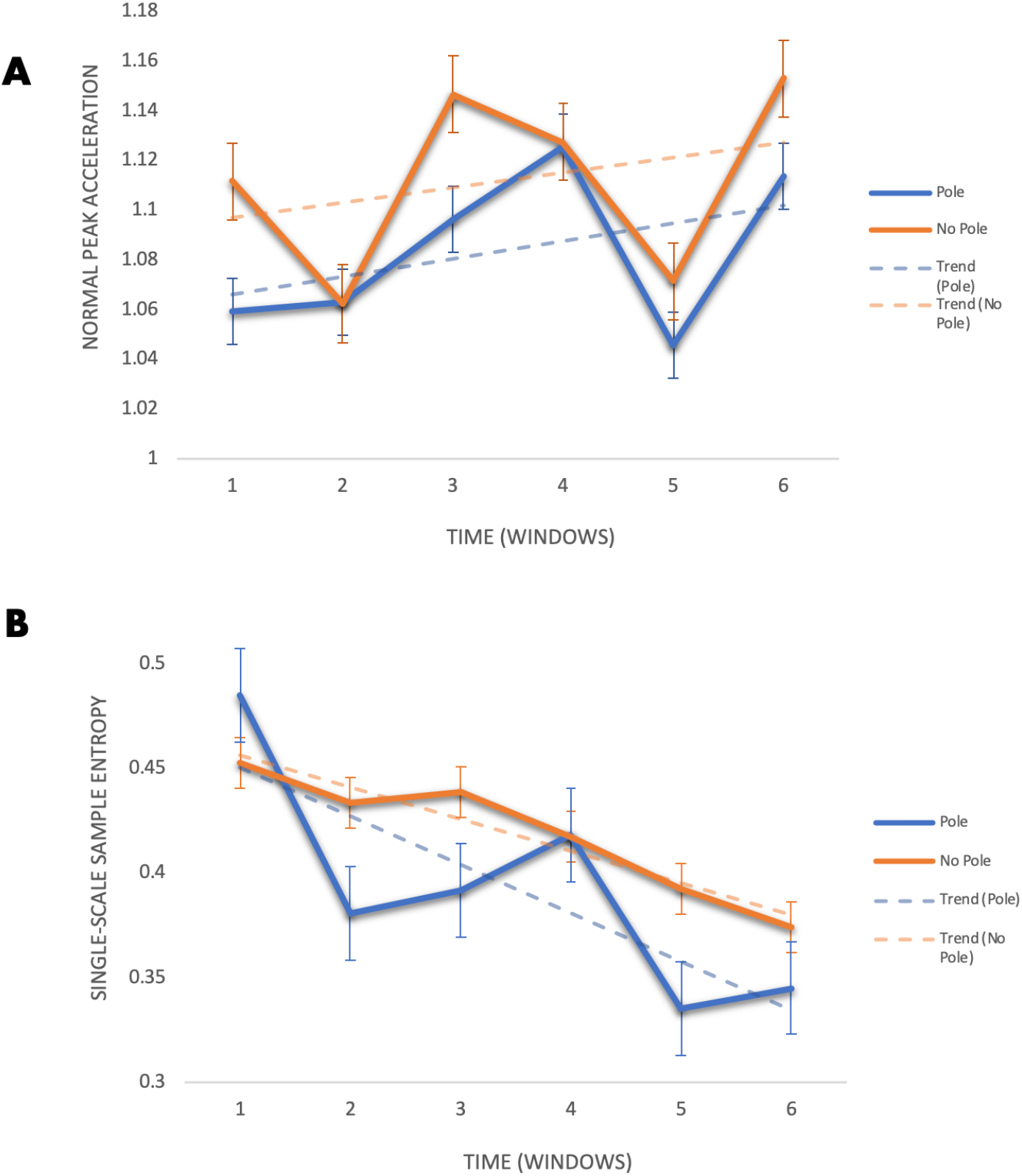
Normal Peak Acceleration (NPA) and Sample Entropy (SampEn). **(A)** NPA Analysis. Average values of NPA per windows 1 to 6 including standard errors. Solid blue line plots average NPA values for plants in pole condition and dashed blue line plots its linear trend. Solid orange line plots mean NPA values for plants in no-pole condition and dashed orange line plots its linear trend. **(B)** SampEn Analysis. Average SampEn values per windows 1 to 6 and standard errors for *m* = 2, *r* = .25. Solid blue line plots average SampEn values for plants in pole condition and dashed blue line plots its linear trend. Solid orange line plots average SampEn values for plants in no-pole condition and dashed orange line plots its linear trend.

Linear mixed effect analysis examined the fixed effect of pole (No Pole = 0, Pole = 1), time (Window = 1 to 6), and their interaction on NPA. The full model also included the random effects of individual plant, plant pair, and the nested random effect of time (Window) on NPA. Examination of changes in model fit (−2LL) with step-wise removal of fixed effects revealed no effect of the pole x window interaction (χ^2^ (1) = 0, p = .99), of the pole (χ^2^ (1) = 1.4649, p = .23), or of time (χ^2^ (1) = 1.402, p = .24) on NPA. The average changes in NPA values suggest that the patterns of variation in the kinematics of plant nutation are more complicated than the gradual shift from more circular to more elliptical shapes commonly reported in the literature^[7, 11, 23]^ (see Figure 3A). Importantly for present purposes, the changes in kinematics identified by NPA do not capture processes that guide plants towards the pole.

### Predictability of Nutation Patterns: Sample Entropy

Sample Entropy (SampEn) analyses provided measures of the predictability (or regularity) of nutation patterns over time. SampEn constructs vectors of length *m* out of the data points and then computes the conditional probability of finding matching vectors of length *m* + 1 within a tolerance radius *r*, if matching vectors of length *m* have been already found within that tolerance radius *r*. Lower SampEn values (values closer to 0) indicate more predictability. Figure 3B displays the mean (and standard error) of SampEn over time for the two conditions when the length of vectors is *m* = 2 and the tolerance radius is *r* = .25. Observed changes in SampEn values in Figure 3B suggest that nutation patterns become more predicatable in time for both conditions and that nutation is more predictable in the presence of the pole.

To test this observation, we run a linear mixed effect analysis to examine the fixed effect of pole (No Pole = 0, Pole = 1), time (Window = 1 to 6), and their interaction on SampEn. The full model also included random effects of plant pair and the nested effect of time (Window) on SampEn. Examination of changes in model fit (−2LL) with step-wise removal of fixed effects revealed a significant effect of time (χ^2^ (1) = 14.973, p < .001) and a significant effect pole (χ^2^ (1) = 4.9049, p < .03) on SampEn (see Table 2).^1^ The interaction between pole and time was not significant. This trend suggests a progressive increase in predictability of nutation patterns as plants grow, and an influence of the presence of a pole in the vicinity of plants in the overall predictability of the time series. Such an influence is a first indicative of the possible effect of the availability of a support to climb on the control strategy of climbing beans’ nutation.

**Table 2.**
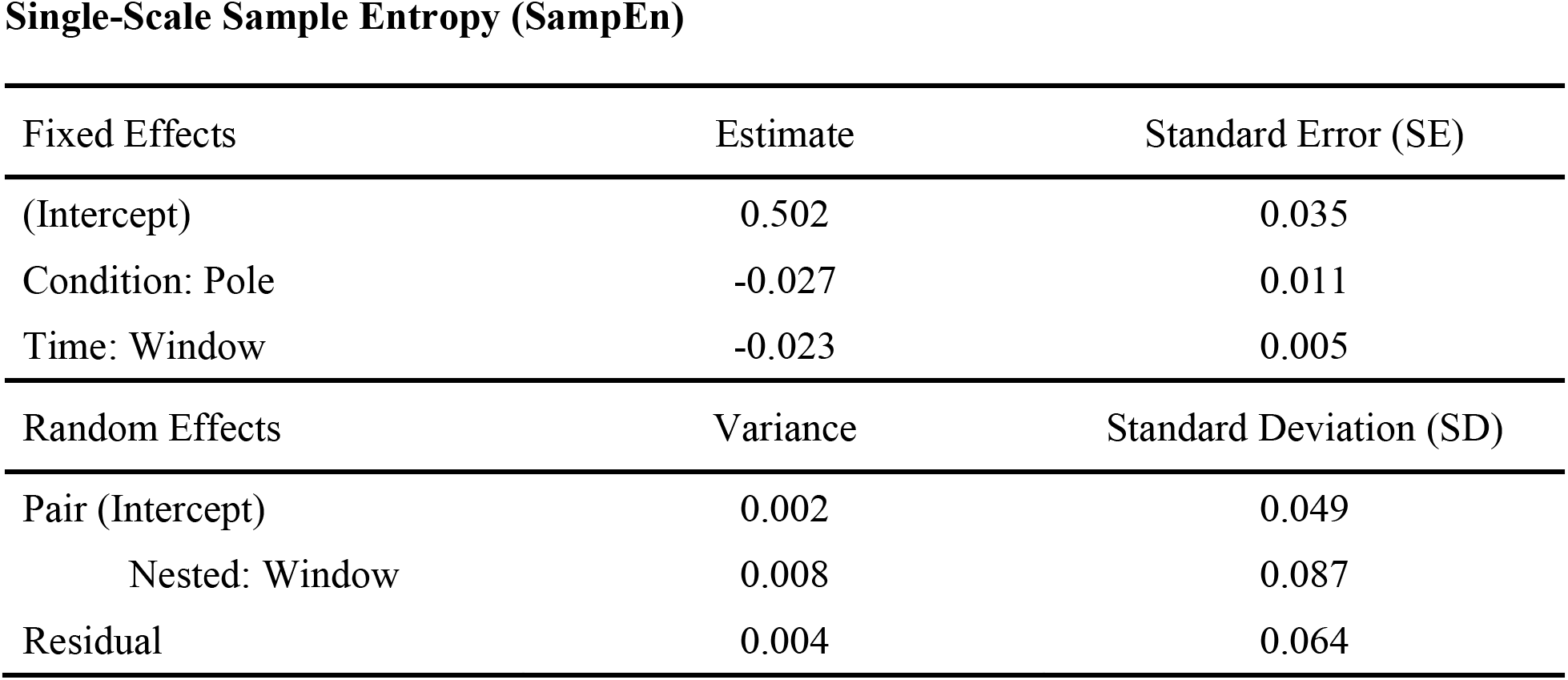
Final model estimated to test the effects of time (Window) and condition (Pole) on SampEn. The model includes all significant fixed effects and all random and nested random effects. Intercept provides an estimate of performance in the base line condition (No Pole = 0). Estimates are the non-standardized weights or coefficients in the linear mixed effect model.

| Fixed Effects | Estimate | Standard Error (SE) |
| --- | --- | --- |
| (Intercept) | 0.502 | 0.035 |
| Condition: Pole | -0.027 | 0.011 |
| Time: Window | -0.023 | 0.005 |
| Random Effects | Variance | Standard Deviation (SD) |
| Pair (Intercept) | 0.002 | 0.049 |
| Nested: Window | 0.008 | 0.087 |
| Residual | 0.004 | 0.064 |

### Complexity of Nutation Patterns: EMD-Based Multiscale Sample Entropy

Multiscale Sample Entropy (MSE) was used to analyze the complexity of nutation patterns, that is, the degree of interactivity of multiscale processes giving rise to nutation. Accordingly, MSE computes SampEn of nutation patterns over time at different temporal scales of the movement. These scales are defined in terms of its intrinsic mode frequencies (IMFs) by a procedure of fine-to-coarse Empirical Mode Decomposition (EMD; see Figure 4A). Higher irregularity (or higher magnitudes of SampEn) on a single scale can indicate less deterministic structure or a greater degree of randomness in the pattern. Higher irregularity that is preserved across two or more temporal scales, however, suggests greater complexity: a mix of interconnected deterministic and stochastic processes evolving across scales typical of adaptive, biological movement systems. This is usually reflected in higher and more stable SampEn values across those scales although the SampEn value of the first scale—the whole signal—may be higher for the least complex data^[53 – 55, 60 – 63]^. Consistent with MSE literature, average SampEn values across scales of analysis in Figure 4B suggest that the irregularity of nutation patterns is slightly higher for the no pole condition at scale 1 (left chart). The irregularity of nutation patterns and its associated SampEn values progressively become higher and more stable in subsequent scales for the plants in the pole condition for at least some windows of time (right chart).

**Figure 4.**
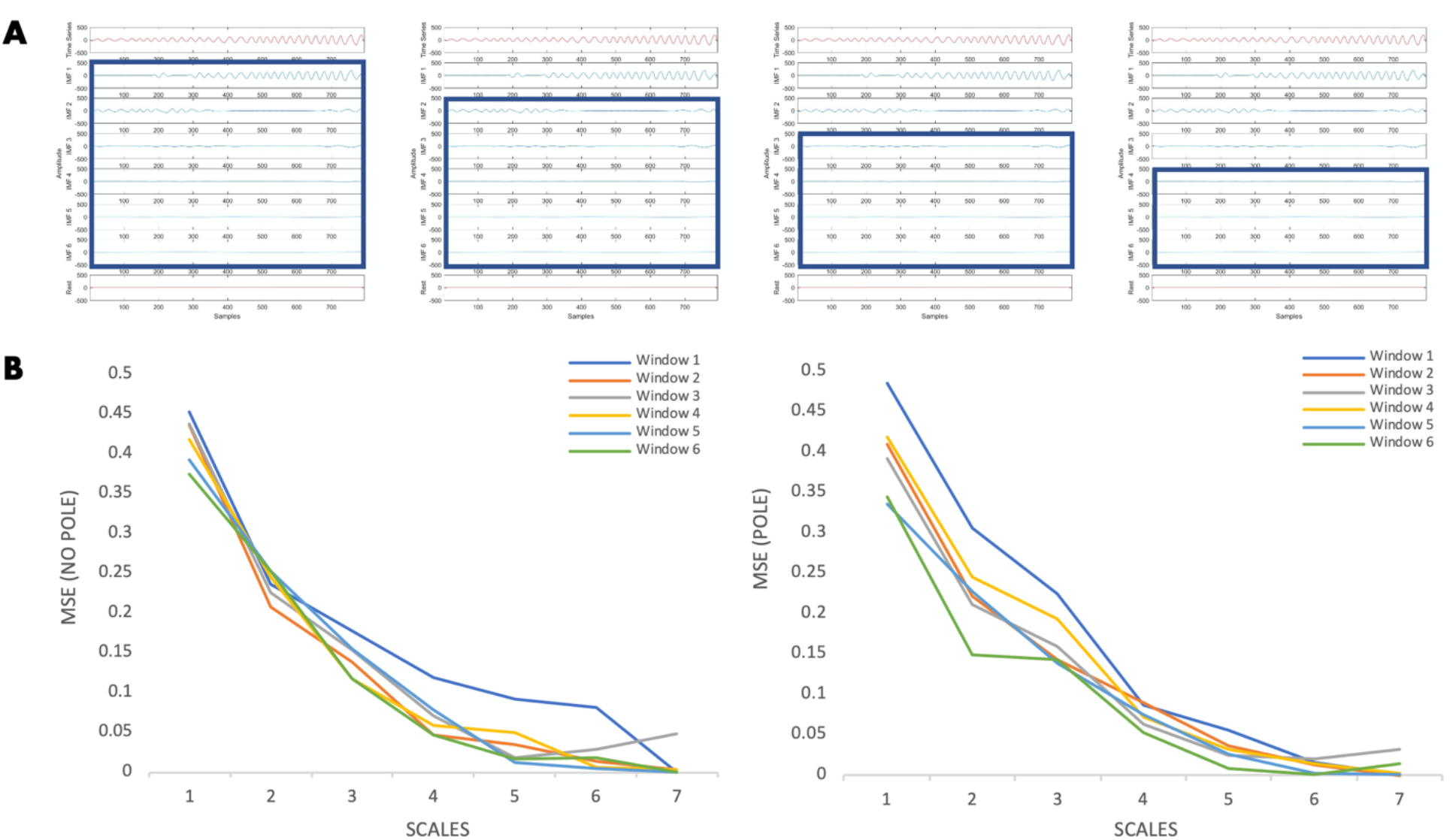
Empirical Mode Decomposition and Multiscale Sample Entropy. **(A)** Example of EMD analysis of a time series and of the fine-to-coarse EMD procedure. The 4 panels show the EMD for the Pair 3, pole condition plant (see Table 1). From top to bottom, each panel shows: full time series (red), six IMFs (blue), and Rest of signal (red). Thick-lined blue squares show IMFs that are included in each Scale generated by a fine-to-coarse EMD procedure. Left panel represents Scale 1 including IMFs 1 to 6 (the whole signal). Second to left panel represents Scale 2 including IMFs 2 to 6 after subtracting the highest frequency component (IMF 1). Second to right panel represents Scale 3 including IMFs 3 to 6 after subtracting the two highest frequency components (IMFs 1 and 2). Right panel represents Scale 4 including IMFs 4 to 6 after subtracting the three highest frequency components (IMFs 1, 2, and 3). **(B)** EMD-based MSE (*m* = 2, *r* = .25) for all windows and scales. Left chart shows no-pole condition. Right chart shows pole condition. General reduction of the average magnitude of SampEn is observed for both conditions. The average magnitude of SampEn is preserved across scales 2 and 3 for windows 3, 4, and 6 in pole condition.

To test this observation, we ran a linear mixed effect analysis to examine the fixed effects of pole (No Pole = 0, Pole = 1), time (Window = 1 to 6), and temporal scales of analysis (Scale = 1 to 7), in addition to all their interactions on SampEn (*m* = 2, *r* = .25). The full model also included the random effect of plant pair along with a nested random effect of time (Window) and the random effect of individual plants along whit a nested random effect of scale on SampEn. Removing the pole x window x scale interaction from the model resulted in a significant reduction in fit (−2LL), χ^2^ (1) = 5.6645, p = .017 (see Table 3, Full Model).^2^ A detailed visual analysis of the average changes in SampEn across scales for the different windows in the two conditions (Figure 4B) guided initial interpretation of this statistically significant three-way interaction. In particular, we observe that the overall reduction in the magnitude of SampEn (i.e., the amount of irregularity in nutation patterns) as we move across temporal scales of analysis follows two different arrangements. Notably, in the no pole condition the magnitude of SampEn is not preserved across any of the temporal scales of analysis. In contrast, when the pole is present the magnitude of SampEn is preserved across scales 2 and 3 for windows 3, 4, and 6. This is a signature of the complex structure in the variability of those patterns at those stages of the movement (Windows = 3, 4, and 6) and at temporal scales (Scale 2 ~ 0.1Hz, and Scale 3 ~ 0.05Hz) that capture a relevant part of the frequency power of the signal (see Figure 4A above).

**Table 3.** Final models estimated to test the effects of time (Window) and condition (Pole) on SampEn at the different temporal scales of analysis generated by a fine-to-coarse EMD procedure (Scale). ***Left panel:*** Full model including all windows and scales. ***Right panel:*** focused model including windows 3, 4, and 6, and scales 2 and 3. Both models include all significant higher-order interactions, all lower-order interactions, all component lower-order effects, and all random and nested random effects. Interactions are marked by ‘*’. Intercept provides an estimate of performance in the base line condition (No pole = 0). Estimates are the non-standardized weights or coefficients in the linear mixed effect model.

| *Full Model* |  | *Focused Model* |  |
| --- | --- | --- | --- |
| Fixed Effects | Estimate (SE) | Fixed Effects | Estimate (SE) |
| (Intercept) | 0.436 (0.029) | (Intercept) | 0.501 (0.074) |
| Condition: Pole | 0.083 (0.028) | Condition: Pole | -0.188 (0.09) |
| Time: Window | -0.007 (0.005) | Time: Window | -0.009 (0.008) |
| Scale | -0.08 (0.009) | Scale | -0.111 (0.025) |
| Pole*Window | -0.026 (0.007) | Pole*Scale | 0.075 (0.035) |
| Pole*Scale | -0.009 (0.009) |  |  |
| Window*Scale | 0.001 (0.001) |  |  |
| Pole*Window*Scale | 0.004 (0.002) |  |  |
| Random Effects | Variance (SD) | Random Effects | Variance (SD) |
| Pair (Intercept) | 0.005 (0.068) | Pair (Intercept) | 0.0004 (0.02) |
| Nested: Window | 0.0004 (0.02) | Nested: Window | 0.001 (0.03) |
| Plant (Intercept) | 0.00009 (0.009) | Residual | 0.009 (0.097) |
| Nested: Scale | 0.0005 (0.022) |  |  |
| Residual | 0.004 (0.064) |  |  |

To determine whether these observed arrangements were statistically reliable, we ran a linear mixed effect analysis to examine the changes of SampEn values for the two conditions (No Pole = 0, Pole = 1) focusing in scales 2 and 3 and windows 3, 4, and 6. The model included the random effect of plant pair and the nested effect of time (Window). Removing the pole x scale interaction effect from the model resulted in a significant reduction of fit (−2LL), χ^2^ (1) = 4.3589, p < .04 (see Table 3, Focused Model). Follow-up analyses of variance showed that average SampEn values were significantly reduced from scale 2 to scale 3 in windows 3, 4, and 6 for the no-pole condition (F(1, 4) = 60.69, p = .001), but not for the pole condition (F(1, 4) = 1.28, p =.32), confirming the arrangements observed in figure 4B. Subsequent MSE analysis of the surrogate shuffled time series of the plants in the pole condition showed the vanishing of the arrangement observed in figure 4B and, therefore, offered further support to the presence of a complex structure in the variability of nutation patterns in windows 3, 4, and 6. In conjunction, these results show that when a climbable structure is present in the plants’ environment, the variability in nutation patterns is more complex, that is, structured by a blend of deterministic and random processes. Such results constitute further evidence for the possible effect of the availability of a support to climb on the control strategy of climbing beans’ nutation.

## Discussion

One aim of our study was methodological: for the first time in the literature, we applied nonlinear methods of behavioral analysis to the dynamics of plant nutation. The direct study of the dynamical features of plant nutation provides a way to understand the movement beyond its kinematical characteristics. The study of the kinematics of nutation usually requires a two-dimensional^[7]^ or even a three-dimensional^[13]^ analysis to reach an understanding of the movement that is allegedly able shed light on its underlying mechanism^[11, 13, 18]^. However, we have shown that there are methods to assess dynamical features of plant nutation from a one-dimensional time-series describing the movement over time—as it is commonly acknowledged in the sciences of complexity^[38, 44, 45]^. Indeed, the proposed methodology allows for a direct and complete study of the dynamics of plant nutation without relying on particular kinematic patterns or *ad hoc* modelling criteria (e.g., modeling climbing plants’ grasping of a support in terms of hypothetical animal wrist-like and digit-like—thumb and index—ensembles)^[64]^. It is important to note that our methodology frees hypothesis-testing from zoomorphic biases.

Another aspect of the methodology is that it provides guidelines for plant scientists to process nutation data, and potentially other plant movements, in a reliable and informative way. The methodology (described in detail in the Methods) can be applied to any movement data collected by time-lapse videos from the zenithal point of view. It provides justification and procedures for choosing one dimension of the movement, for assessing the frequency-resolution needed to capture the relevant information, for assessing stationarity, for selecting the windowing of the data, and for selecting the parameters of the entropy analyses. In this way, the proposed methodology becomes a basis for the generalization of nonlinear methods of behavioral analysis for plant movements.

When the methodology was applied to the study of common beans’ nutation in two conditions (pole vs. no pole), the way dynamical measurements allowed for a deeper understanding of the movement became manifest. Our hypothesis was that, if the presence of a support to climb had an effect in the control of nutation, this fact would be reflected in the dynamical properties of the movement. In particular, common beans’ nutation with a support to climb in their surroundings would exhibit common signatures of adaptive, controlled movements in terms of nonlinearity, predictability, and complexity. The results of our study show these three signatures in nutation, strongly suggesting that the support to climb is to be regarded as a relevant environmental factor for the control of nutation.

This result speaks to the mechanism of nutation and its relationship to the kinematics of the movement. First, the results of the analysis of harmonicity show that the commonly reported gradual shift from more circular to more elliptical shapes in the nutation patterns of climbing beans is actually a non-gradual, elaborate shift and is not related to the possible adaptive processes that guide plants towards a support to climb. And second, the differences found both in single-scale and multiscale sample entropy for the two conditions suggest that a mechanism based on endogenous, purely linear oscillators^[2, 13, 16, 18]^ are not good enough to capture the complexity of nutation in different environmental settings. Instead, a mechanism that is not merely controlled by endogenous factors, and that is sensitive to the position of a support to climb in the environment of the plant, proves to be a better candidate.

Last, the complexity shown by nutation patterns in our study when a support to climb was present in the surroundings of the plants is compatible with the finding that complexity in plants’ electrical signaling may be driven by external factors^[67]^. More research is needed to fully understand nutation dynamics and to corroborate both its adaptive, controlled nature—e.g., exploring nutation in other plants, modelling nutation both at the behavioral and cellular scales, etc.—and the seemingly intermittent influence of the support to climb onto. These may be due to other conditions intermittently affecting the movement. Overall, the proposed methodology is a robust tool to pursue this kind of research. With this methodology, plant scientists can improve the understanding of the dynamics of plant movements and gain perspective with regard to the fundamental features and underlying mechanisms of plant nutation.

## Methods

### Plant Materials

Experiments were performed in the common bean (*Phaseolus vulgaris* L.). Seeds of the cultivar “Buenos Aires” were provided by Semillas Ramiro Arnedo S.A., Spain <http://www.ramiroarnedo.com>.

### Germination and growth conditions

Plants were light-grown from seed in a controlled-environment chamber (300 × 400 × 240 cm.) at the Minimal Intelligence Lab (MINT Lab), University of Murcia, Spain.

Prior to sowing, Buenos Aires var. common bean seeds were germinated on filter paper, soaked in an aqueous solution of hydrogen peroxide (H_2_O_2_)—15mL distilled water/3mL H_2_O_2_— to promote germination in Petri dishes kept in darkness for 24-48 h. Upon germination, young seedlings were transferred to coconut fiber growing pellets and kept on propagation trays. Humidity levels were checked periodically to maintain a moist bedding.

Once the first pair of true leaves had appeared, healthy-looking seedlings were transplanted into a mixture of peat moss and perlite (70%-30%) in the center of small black plastic pots (70×70×80mm. in depth). The temperature of the growth chamber was kept at 23°C (day) and 19°C (night) ± 1 °C, photoperiod at L16:D8 h. light cycle, and relative humidity 55% ± 5%. Light was delivered from the top (photosynthetically active radiation (PAR) photon fluence rate of 235 ± 25 μmol m–2 s–1 at leaf level—Delta OHM HD 9021 solid state PAR sensor, Caselle di Selvazzano (PD), Italy).

Plants were watered (filtered tap) two hours into the photoperiod. All seedlings received the same amount of water per day; amount that progressively increased with plant age from 20 to 60mL. No other treatment was applied.

### Stimulus and experimental design

Measurements were carried out from the onset of ‘bending’: a morphological feature characteristically displayed in *P. vulgaris* as the apical segment below the horizontal section of the shoot apex in ca. two-week-old plants becomes curved^[11]^.

Plants were pairwise compared with an eye to selecting the most similar seedlings as model plant pairs, prior to transfer to the recording booths. Pairwise comparisons were based on uniformity in stem and internode length and diameter, leaf shape and total leaf area. The most uniform subjects that were ready to be transferred into the recording booths were selected. These were transplanted to larger pots (Air Max 7 liters square plant pots—200×200×270mm. in depth) filled with the same mix (70% peat moss/30% perlite). Potted plants were saturated with filtered tap water, and allowed to drain, to avoid any further watering during time-lapse recordings. Pilot tests^[27]^ confirmed that humidity levels could be kept within an optimal range for the duration of the experiment.

Pairs of potted plants were then randomly allocated to one of two cylindrical booths within the growing chamber, where they were placed at the center of the bottom cylinder. Booths were identical with the exception of a vertical pole (height: 0.90 cm; diameter: 1.8 cm.) presented as a stimulus—a potential support for the bean to twine around—in only one of the two booths. The pole was placed at a distance of 30 centimeters from the plant center. Before starting recordings, plants were allowed to stabilize for two hours after having been transferred.

Each booth consisted of a right circular cylinder with a radius of 93 cm. A height of two meters of the vertical cylindrical surfaces, together with the disposition of the booths within the growth chamber, was such that it prevented air circulation produced by the chamber’s extractor (RVK 1000m3/h) and injector (PK150-L 780m3/h) used for air renewal and temperature regulation from affecting plants’ nutation. To distribute light more uniformly, the cylindrical surfaces were covered in white reflective film.

The light source consisted of a high-pressure sodium lamp (Lumatek pulse-start HPS Lamp 250W) with specific horticultural gas blend for optimal spectral output consistency and PAR/PPF level maintenance for plant growth. Symmetrical lighting was ensured by suspending lamps vertically within the cylindrical booths under a white parabolic reflector. Lamps were positioned centered at 150 centimeters above each potted plant, providing a photon fluence rate of 430 ± 50 μmol m–2 s–1 at leaf level.

### Video recording

Time-lapse cameras (Brinno TLC200 PRO) were placed within each booth 130 cm. off the ground, pointing vertically down to the center of the potted plant. Time-lapse records were made, the time interval between frames for data collection of the position of the shoot tip at one frame per minute. A 4.2 μm High Dynamic Range (115dB) image sensor made time-lapse recording possible during the 8 hours of darkness under a dim phototropically inactive green safelight with a fluence rate under 5 μmol m–2 s–1. The bottom face of the cylinder was covered in black to maximize the contrast between the shoot tip and the background whilst recording.

### Data Processing

The first processing step is to digitize the 2D coordinates of shoot tips obtained from time-lapse videos. This procedure introduces noise in the time-series, which can be minimized if the selected level of resolution is not too high considering the rate of change in the position of shoot tips. To that end, we digitized the time-lapse videos using resolutions of 24 Hz, 6 Hz, and 3 Hz, and computed the spectral properties of the resulting three time-series using Fast-Fourier transformations. The transformations showed that the fundamental frequencies of the movement were concentrated under 0.5Hz (M ± SD = 0.29 ± 0.05; Figure 2A above) for the three tested frequencies, ensuring the robustness of our digitalization process. We subsequently selected the central resolution value, 6 Hz. This choice guarantees that the resulting 2-D time-series of the plants’ shoot tips includes the relevant information about the nutation movement while avoiding meaningless noise. It also guarantees long enough time series for dynamical analysis.

In line with the literature on dynamics, we assume that the underlying dynamical organization of biological rhythmic movements can be appropriately reconstructed from a single measurement of the behavior of interest^[45]^. Thus, for all the analyses described below, we have only used the digitalized position time-series of the plants’ shoot-tips in the horizontal axis of the video frames.

### Stationarity Testing

Two of the three dynamical analyses used in the present study (*Harmonicity* and *Single-Scale Sample Entropy*) assume stationarity of time-series data. To test this assumption, we initially applied a Dickey-Fuller test (ADF Test)^[51]^. This test examines the null hypothesis that a unit root—i.e., a stochastic or unpredictable trend or random walk drift in a time-series—is present in a time series sample. If a unit root is not found in a time series, there are no grounds for rejecting the null hypothesis, and stationarity is concluded^[50]^. We selected this test because of its clear interpretative cut-off to distinguish between stationary and non-stationary data. However, the ADF test suffers from low statistical power and can only exclude strong violations. As a countermeasure to this limitation, we employed a two-step procedure to determine whether weaker violations of stationarity were at play or not^[63]^. The first step consists in applying transformations to the data to *make them* stationary. These transformations involve either differencing the time-series or fitting a polynomial function and subtracting the fitted trend from the data. The latter transformation is known as detrending or de-seasoning, depending on the way the polynomial function is subtracted from the original time series^[48, 49]^. The second step consists in applying the analyses that assume non-stationarity to both the original and the transformed data, and comparing the results. If results are indifferent to the transformation, it is further evidence that the assumption of stationarity is not being violated. These methods, however, entail a manipulation of the data that might cause the discarding of important information of the time series, so they must be used carefully.

### Window Selection

To examine whether the effect of the pole on nutation dynamics change over time, we parsed the data into time-windows. We used a method for window selection that accomplishes two critical goals. First, each window must include enough data points to reveal the underlying dynamical organization of nutation patterns. A good rule of thumb for the analyses employed here is a minimum of 100 data points. And second, windows must not include many overt variations in amplitude and frequency that can impact the results of the dynamical analysis. Accordingly, we selected hoping windows of 100 data points, starting from the last point defined by the moment at which the plant in the pole condition touched the pole. For most plants, this procedure guaranteed at least 6 windows of 100 points that could be used for the analysis. These relatively small windows prevented large variance in the amplitude and frequency of the time series within the windows (helping ensure stationarity), while providing enough data points for the dynamical analyses.

### Data Analysis

Measures of harmonicity (Normal Peak Acceleration; *NPA* hereafter), regularity (Single-Scale Sample Entropy; *SampEn* hereafter), and complexity (Multiscale Sample Entropy; *MSE* hereafter) of nutation patterns were computed for each plant in each of the six analyzed windows. The analyses performed to compute each of these measures will be described in turn.

#### NPA

The Fast-Fourier transformation (FFT) of the position time-series performed during *Data Processing* (see above) indicated that 99% of the power in the nutation signals was under 1Hz. Thus, to minimize the influence of noise in the computation of *NPA*, all time-series were filtered using a dual-pass second-order Butterworth filter with a cut-off frequency of 1.5 Hz. After filtering, the harmonicity of nutation patterns was then computed based on the average of the peak acceleration values obtained for each half-cycle^[52]^. For a purely sinusoidal (or harmonic) movement of amplitude A and frequency ω, peak acceleration is equal to Aω^2^. *NPA* was computed by dividing the observed peak acceleration by Aω^2^ and is, thus, a sensitive measure of the degree of harmonicity. A *NPA* of 1 is a signature of harmonic patterns, consistent with purely linear control processes. Deviations of *NPA* from 1 index a departure from a purely harmonic pattern, signaling contributions of non-linear processes to the movement of nutation.

#### SampEn

Generally speaking, different entropy measurements of time series (e.g., Kolmogorov–Sinai entropy, approximate entropy, *SampEn*) aim to assess the probability of knowing future states of the series given measurements of their current states: the more predictable (or regular) the time series is, the lower entropy it has, and vice versa^[47, 56]^. In the case of *SampEn*, such an assessment is pursued in a way inspired by approximate entropy^[57]^. The general method of *SampEn* consists of the comparison of vectors of different length, *m* (e.g., m = 1, m = 2, m = 3, and so on), given a tolerance for assessing similarity, *r*. For example, vectors *m* = 2 or *m* = 3 are those composed of 2 or 3 data points of the time series respectively. And a tolerance of *r* = 0.25 defines how much different two vectors can be so as to be deemed similar to one another. The method of *SampEn* calculates the conditional probability of finding a match for a vector of *m* + 1 data points given that it has already found a match for a particular (template) vector of *m* points^[44, 53]^ within a tolerance radio *r*.

In terms of its two parameters, literature on *SampEn* recommends vector lengths of *m =* 1 or 2 and tolerance radiuses, *r*, between 0.1 and 0.25 of the standard deviation of the time series^[44]^. However, given the properties of the time series of plant nutation, especially its temporal scale (i.e., small displacements from data point to data point) and the general homogeneity of the movement, it makes sense to consider longer vectors (e.g., *m* = 3) than those considered in faster, more variable human/animal movements. Also, different data driven methods of parameter selection have been recently proposed like, for example, the calculation of parameters that relies on autoregression functions of time series^[58]^. Another option is to estimate the *m* value of a time series by plotting its median *SampEn* values as a function of different values of *r* and selecting the *m* values at which the curves become more similar^[59]^. In the case of parameter *r*, the relative error of *SampEn* is calculated and plotted against *r* values^[59]^. The *r* value at which relative error is minimal should be chosen for the analysis. We employed these methods for parameter selection in this study and results suggest that, as a general rule, *m* = 2 or 3 and *r* = 0.25 are reasonable selections.

#### MSE

Higher values of *SampEn* (i.e., a measure of reduced regularity) have been interpreted to indicate greater complexity in movement patterns^[58, 59, 63]^. However, this is problematic because its value is maximized (around 2) for completely random, unstructured patterns. Such a lack of correspondence may lead to situations in which less complex time series get higher *SampEn* values than more complex ones^[60 – 62]^. Complex patterns are characterized by structured variations (i.e. subtle time correlations) in the midst of randomness, reflecting the adaptability (or context sensitivity) of biological patterns^[39, 68 – 71]^. Importantly, this rich mix of order (regularity) and disorder is preserved across scales when the pattern is complex^[54, 60, 61, 72 – 74]^.

*MSE* assesses the degree of disorder (*SampEn*) in multiple scales and allows assessment of the extent to which it is preserved across scales. Thus, a *MSE* analysis provides a way to disambiguate between random and complex patterns ^[39, 60]^. In particular, at the more microscopic scale, the non-complex time series (e.g., white noise) have a higher *SampEn* value. However, more complex time series exhibit higher and, more importantly, more stable values of *SampEn* at progressively more macroscopic scales (at least in the relevant scales where structure exists). For non-complex pattern, an exponential decrease in *SampEn* is observed as the scale of analysis is increased.

Performing *MSE* analyses requires making a decision regarding the way the different spatiotemporal scales are defined. A “coarse-graining” method in which different scales are generated by averaging the data points of the time series within non-overlapping windows of increasing length—e.g., averaging data points in groups of 2 points, 3 points, 4 points, etc.—has been proposed^[39, 60]^. However, this method has the problem of shortening the time series to a half, to a third, to a quarter, and so on, as the windows of data points increase in length. For this reason, it is not adequate in the case of regular or short time series. We employed instead an empirical mode decomposition analysis (*EMD*) to carve out the relevant spatiotemporal scales of the time series of nutation^[55]^.

*EMD* is a data driven methodology that consists in finding the intrinsic frequency modes (*IFM*) of time series and combining them to generate the different scales. Once these modes are found, there are two ways to perform an *EMD*: (i) coarse-to-fine *EMD*, in which the lowest-frequency *IMF* is subtracted from the signal in each step until only one *IMF* is left; and (ii) fine-to-coarse *EMD*, in which the highest-frequency *IMF* is subtracted from the signal in each step until only one *IMF* is left. Usually, most of the power of the signal in time-series of plant nutation is in the highest-frequency ones, so a fine-to-coarse *EMD* is recommended to get a better understanding of the significant differences between scales.

Finally, to confirm that the complexity observed in *MSE* analyses was not due to an artifact of the analysis, we compared the *MSE* analysis of original time series with the *MSE* analysis of their corresponding surrogate shuffled time series. Shuffling (i.e., randomly reordering) the original time series destroys their temporal dependencies while preserving their statistical features. Therefore, surrogate shuffled time series are by definition less complex than the original time series and the features associated with complexity (e.g., the stability of *SampEn* values across relevant scales) in their *MSE* analysis must vanish if they are not an artifact.

### Statistical Analysis

We used the *lme4* package^[65]^ of the *R Studio*^[66]^ suite to estimate linear mixed effect regression models to examine the contributions of the pole (No pole = 0, Pole = 1), time (Windows 1 to 6), and their interaction to three outcome measures computed to characterize three aspects of plant dynamics: non-linearity, predictability (regularity), and complexity. When modeling complexity, we additionally examined the effect of the temporal scale of analysis (Scales 1 to 7) and its interactions with pole and time. In all cases, we started by identifying the random effects of plant pairs and of the individual plants that should be included in the model. Null models with different combinations of random effects were systematically estimated and the most conservative, best fitting model that converged was selected. We then implemented a backwards approach to identify statistically meaningful fixed effects. The full model included the fixed effects of time, pole, and scale (when pertinent), and their interactions in addition to the random effects previously identified. Models were trimmed by removing fixed effects individually, starting from the higher-order interaction effect. At each step, we compared the deviance (−2 Log Likelihood; −2LL hereafter) between a larger model and a simpler nested model that excluded the predictor under analysis. The change in −2LL follows a chi-square distribution with degrees of freedom equal to the difference in the number of parameters between nested models, allowing for a test of statistical significance of each fixed effect. Final models included only significant higher-order interactions, all lower-order interactions, and all component lower-order effects.

## Footnotes

1 Similar effect of window on SampEn, but not of pole, was found when vector length is *m* = 3.

2 A similar marginal effect of the pole x window x scale interaction on SampEn was found when the vector length is *m* = 3 (p = .07).

